# Re-analysis of CLAP data affirms PRC2 as an RNA binding protein

**DOI:** 10.1101/2024.09.19.613009

**Authors:** YongWoo Lee, Priyojit Das, Barry Kesner, Michael Rosenberg, Roy Blum, Jeannie T Lee

## Abstract

Using CLAP methodology, Guo et al. recently concluded that PRC2 is not an RNA binding protein (RBP). They suggest that prior findings were CLIP artifacts and argue against RNA’s direct role in PRC2 regulation. Here, we re-analyze their raw datasets and reach contrary conclusions. Through an independent computational pipeline, we observe significant PRC2 enrichment throughout the transcriptome, including XIST. Applying the authors’ published computational pipeline also reaffirms PRC2 as an RBP. Detailed investigation of the authors’ pipeline reveals several unconventional practices. First, Guo et al. retained reads from foreign species and other unmappable reads to obtain a normalization factor. Second, they selectively removed read duplicates from the mappable fraction, while retaining them in the unmappable fraction. Finally, the authors applied an arbitrary cutoff for enrichment values in XIST. Their pipeline thereby inflated PRC2’s background reads and suppressed mappable signals, creating the impression that PRC2 is not a robust RBP.

## INTRODUCTION

Many studies have shown that Polycomb repressive complex 2 (PRC2) is an RNA binding protein (RBP) that associates with a large family of transcripts in mammalian cells. Affinity capture methods such as native RNA immunoprecipitation (RIP) ^1–3^, UV-crosslinked immunoprecipitation (CLIP) ^4–6^, and denaturing CLIP ^7^ have identified large RNA interactomes inclusive of thousands of transcripts in vivo ^2,5–7^. In affinity capture studies that use an RNA bait to survey interacting proteins, PRC2 subunits have turned up as direct interactors for transcripts such as XIst and TERRA RNA ^8,9^. Furthermore, RNA crosslinked proteomic surveys have identified PRC2 components as interacting partners ^10^. Studies show that RNA contacts PRC2 along dispersed amino acid patches on its surface ^11–14^ and binds the EZH2 subunit near its catalytic domain ^14^. Subunits JARID2 ^14–16^, AEBP2, and RBBP4 ^8^ also contact RNA. The PRC2 subunits are now known to recognize RNA via several motifs, including CU-rich and G-rich sequences ^7,17,18^. Some of the G-rich sequences fold into RNA G-quadruplexes (rG4) in vivo, bind PRC2 with 10-90 nM affinities ^19^, and function in PRC2 recruitment ^20,21^, regulation of EZH2’s catalytic activity (H3-K27 trimethylation) ^2^ ^19^ ^4,15–18^, and eviction from chromatin^17^. They have also been implicated in POL-II transcriptional pausing at active genes ^7,22^ and induction of B2/ALU ribozyme activity during the stress response ^23^ ^24^.

Given the existing body of work, the recent work of Guo et al. claiming that PRC2 is not an RBP ^25^ is perplexing. The authors state that prior conclusions about PRC2’s RNA binding capacity ^1,6,7,14,15,17,26,27^ stemmed from reassociation artifacts during RIP and CLIP procedures. RIP and CLIP depend on antibody-based precipitation of the protein of interest, which in turn would pull down interacting transcripts, if they are bound inside cells. The low nanomolar affinities of antibody-antigen binding are known to constrain the stringency of washes during the purification step and could, in theory, lead to reassociation artifacts — i.e., non-interacting RNAs binding nonspecifically to PRC2 during cell lysis, immunoprecipitation, and/or RNA purification. These nonspecific RNAs would then be captured during library preparation and lead to false positive results. Such limitations have spawned variants of the CLIP method, such as denaturing CLIP (dCLIP) and iCLIP ^7,28^. In the latest iteration, Guo et al. developed CLAP (Covalent Linkage and Affinity Purification) and provided evidence for its superior performance relative to CLIP ^25^. CLAP employs Halo-tags for crosslinking PRC2 components to substrate-coated beads and uses denaturing washes to remove non-crosslinked co-precipitants ^25^, which would thereby minimize reassociation artifacts. Through the lens of the authors’ analytical approach and interpretation, CLIP replicated the large RNA interactome for PRC2 but CLAP failed to identify any enriched RNA, including Xist/XIST. The authors therefore urged a critical re-evaluation of RNA’s role in regulating PRC2 activity.

In light of the conflicting publication, we re-analyze the raw datasets deposited in GEO by Guo et al. and ask whether their analysis pipeline contributed to the observed differences. To our surprise, our re-analysis of the same raw data — using the authors’ computational method as well as our own independent pipeline — produced significantly different outcomes from their published account. Indeed, our analysis indicates that CLAP reaffirms the longstanding notion that PRC2 is an RBP. Thousands of transcripts, including the previously identified XIST RNA, exhibit significant enrichment over their corresponding input levels for binding to various PRC2 subunits. By scrutinizing the computational pipeline of Guo et al., we trace the sources of discrepancy to inclusion of unmappable reads and duplicates, and to the application of an arbitrary cutoff for enrichment calculations.

## RESULTS AND DISCUSSION

### Re-analysis using an independent bioinformatic pipeline affirms PRC2 as RBP

Guo et al ^25^. employed cell mixing practices to determine the signal-to-noise ratio in CLIP and CLAP experiments. V5-Halo-tagged proteins of interest were expressed from the plasmid transfection in either mouse or human cells for affinity capture. Plus (+) tag cells harbor tagged proteins of interest, either in human or mouse cells. The +tag cells from one species are then mixed with wildtype cells from the other species prior to performing CLIP/CLAP. When the +tag is expressed in human cells, the admixed wildtype mouse cells provide a control for reassociation artifact in human CLIP/CLAP experiment. Reciprocally, when the +tag is expressed in mouse cells, the admixed -tag human cells provide a control for reassociation artifact in the mouse CLIP/CLAP. In the latter case, if human RNA reads are enriched in the mouse CLIP or CLAP, the human signals must result from reassociation artifacts during the procedure, as the human cells lack the Halo-tagged target protein. In Guo et al., CLIP analysis of PRC2 subunits with the -tag human cells accumulated significant human RNA reads (Fig. 1A, inset, left panel (“G”)), leading the authors to conclude that CLIP is prone to procedural artifacts ^25^. In contrast, the authors reported no such artifacts when performing CLAP ^25^ (Fig. 1A, inset, right panel (“H”)). From these results, the authors argue that CLAP is a superior method to CLIP. Furthermore, the authors’ CLAP showed no enrichment of PRC2 binding in human XIST (Fig. 1B, inset, right panel (“C”)), contrasting sharply with enrichment of SAF-A, PTBP1, and SPEN in various domains of XIST ^25^. The authors therefore conclude that PRC2 is not an RBP.

**Figure 1.**
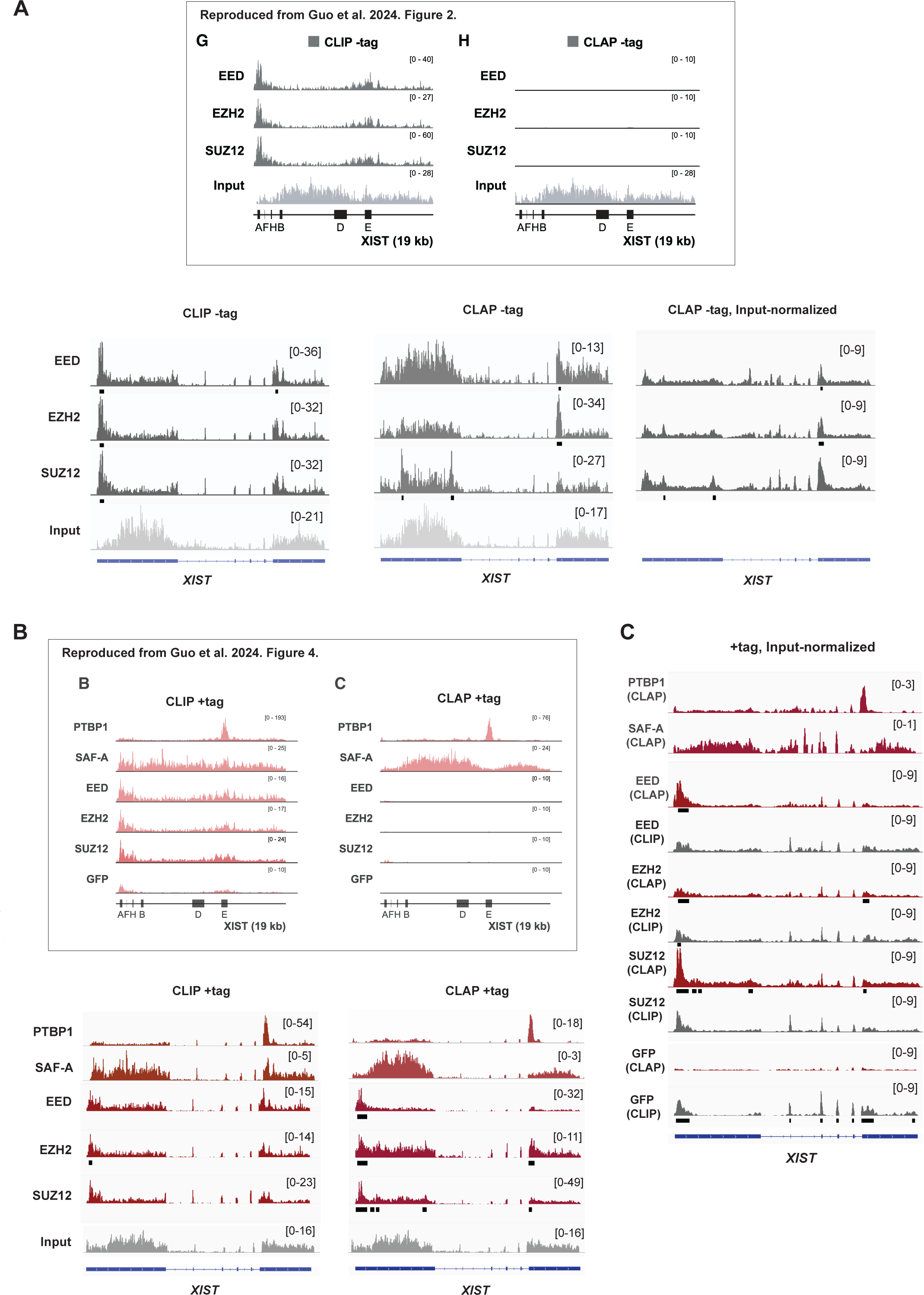
CLAP affirms that PRC2 is an RBP. (A) RNA-binding profiles of PTBP1, SAF-A, and each PRC2 subunit (EED, EZH2, and SUZ12) in -tag human samples, plotted across the human XIST for CLIP and CLAP. Top panel: Spliced XIST transcript (reproduced from Fig.2 G, H of Guo et al. ^25^). Bottom panel: Reanalysis showing unspliced XIST gene. Panels: RPM-normalized RNA reads (Left: CLIP, Center: CLAP); Right: Input-normalized CLAP enrichment profiles. Black bars under each track indicate significant peaks. (B) RNA-binding profiles of PTBP1, SAF-A, and each PRC2 subunits in +tag human samples. Top panel: Spliced human XIST transcript (reproduced from Fig. 4 B, C of Guo et al. ^25^). Bottom panel: Reanalysis showing unspliced XIST gene. Left/Right panels: RPM-normalized RNA reads (CLIP: Left, CLAP: Right). Black bars under each track indicate significant peaks with Y-axis in brackets. (C) Input-normalized enrichment profiles for +tag human samples across XIST. Black bars denote significant peaks.

Given this discrepancy, we re-analyzed the raw sequencing data using our independent computational pipeline. Following widely accepted practices in the field ^29,30^, we removed structural RNA (stRNA), including rRNA, 45S pre-rRNAs, tRNA, snoRNA, and snRNAs, and all duplicates (“de-duplication”) to avoid challenges and artifacts caused by unmappable reads and PCR over-amplification. Our re-analysis of the raw sequencing files revealed both similar and strikingly divergent results. Notably, our re-alignment and enrichment analysis of the authors’ CLIP datasets showed that the profiles for PRC2 subunits, SAF-A, and PTBP1 in human XIST RNA were consistent with these proteins being robust RBPs. These profiles were strikingly similar to the CLIP profiles presented by Guo et al. (Fig. 1B, bottom section) ^25^. Additionally, input-normalized CLAP profiles for PTBP1, SAF-A, and GFP generated by our pipeline were also comparable to those reported by Guo et al. ^25^ (Fig. 1C). Thus, for a majority of samples, our independent analytical pipeline produced enrichment profiles that closely matched those of Guo et al.

However, when we applied the same bioinformatic pipeline to the deposited raw data for PRC2 subunits — EZH2, SUZ12, and EED — both RPM-normalized and input--normalized CLAP profiles for these subunits showed a significant deviation from the results reported by Guo et al. ^25^ (Fig. 1B, bottom section, CLAP +tag, and inset, right panel (“C”; Fig. 1C)). Notably, we observed substantial enrichment in XIST, in sharp contrast to the near-absence of signal reported by the authors ^25^. Remarkably, the CLAP profiles generated by our pipeline closely matched the CLIP profiles for all three PRC2 subunits. This observation indicated that results from the CLIP and CLAP techniques, as applied by the authors, were in fact in close agreement with each other.

To identify significant PRC2 binding regions, we detected statistically significant peaks relative to the reference input using data processed by our pipeline for each protein of interest. In XIST RNA, significant binding was reproducibly observed at the 5’ end around Repeat A (RepA), a motif previously identified as a PRC2 binding site both in vitro and in vivo ^1,15,31^. We note that the +tag enrichment for PRC2 was markedly greater than the -tag “background” (compare Fig. 1A to 1B). In the -tag CLAP negative controls, the 5’ enrichment was notably absent (Fig. 1A). When input-normalized, the enrichment peaks in XIST persisted in the CLAP experiment (Fig. 1C). Thus, our re-analysis of the raw data confirms that the 5’ end of XIST, including Repeat A, serves as a PRC2 binding domain.

Significant binding peaks were detected across the transcriptome, with two notable examples including CTBP1 and ALYRTF (Fig. 2A). The PRC2 +tag CLAP samples consistently showed more prominent peaks compared to the -tag samples (Fig. 2B). Motif analysis of the significant binding sites for EZH2, SUZ12, and EED in the +tag samples identified G-rich motifs for all three subunits (Fig. 2C). This finding is consistent with prior biochemical and in vivo studies highlighting a preference for G-tracts and G-quadruplex-forming sequences ^1,7,17,18^. These results reinforce the concept of sequence-specific in vivo binding of PRC2 in the mammalian transcriptome.

**Figure 2.**
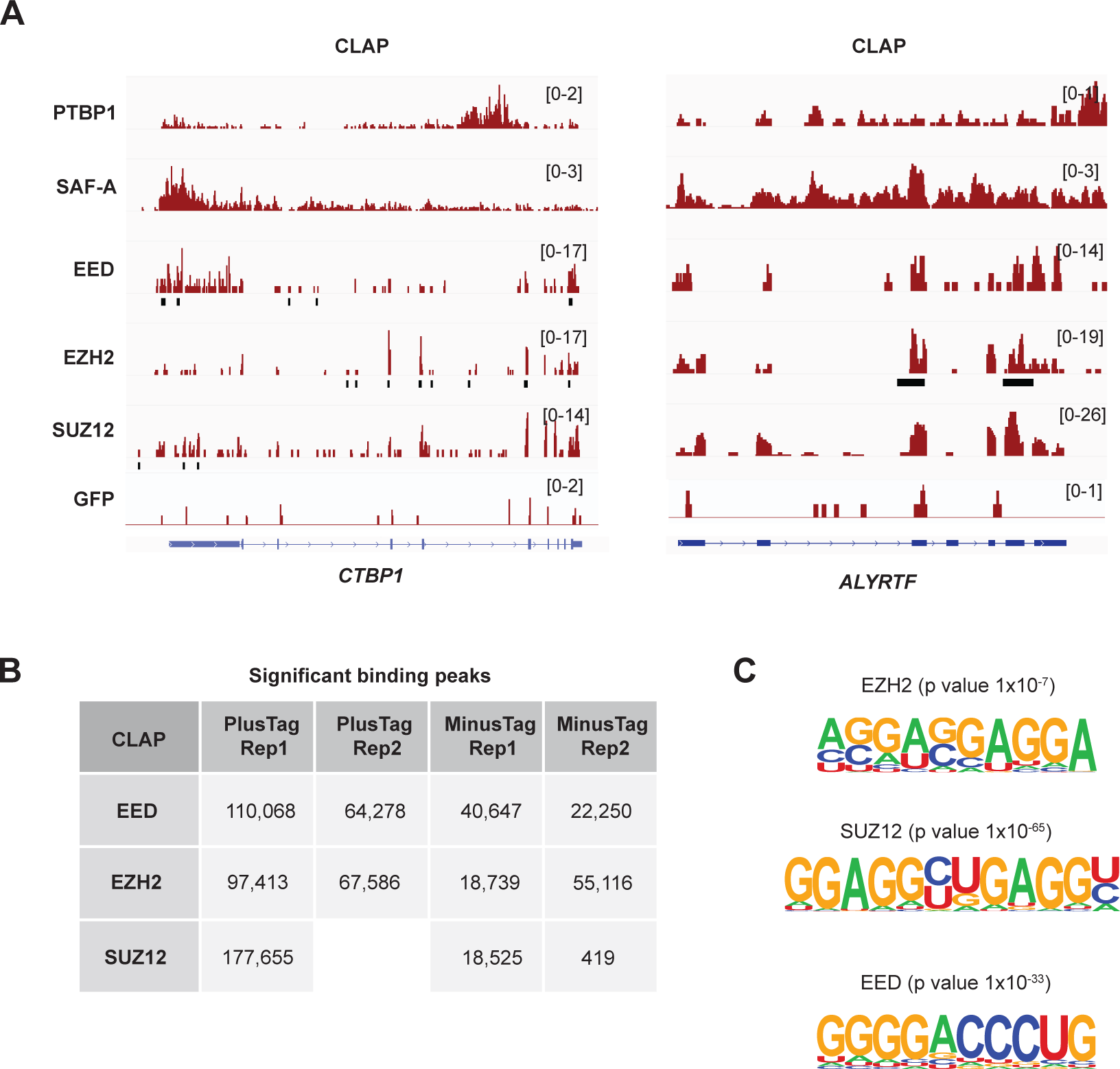
PRC2 binding across the transcriptome. (A) RNA-binding profiles of representative PRC2 target transcripts CTBP1 and ALYRTF (RPM-normalized reads) display significant peaks. (B) Table summarizing counts of significantly enriched over input CLIP and CLAP peaks in the indicated datasets. Blanks indicating missing datasets in Guo et al.’s study. (C) Motif analysis from the +tag PRC2 CLAP experiments (EZH2, SUZ12, and EED).

Guo et al. employed the known RBPs — SAF-A, PTBP1, and SPEN as positive controls to exclude PRC2 as an RBP ^25^. To assess the performance of CLAP for these RNA-binding proteins, we re-applied our informatics pipeline, which was initially used for analyzing PRC2 components. For both SAF-A and PTBP1, re-analysis of CLIP and CLAP datasets showed similar gene alignment patterns (Fig. 3A). Along ELAVL1, for example, CLIP and CLAP demonstrated significantly enriched peaks in identical RNA regions for SAF-A and PTBP1 (Fig. 3A, left panel). Input-normalized tracks also show similar alignment patterns between CLIP and CLAP over BTG2 and CFL1 (Fig. 3A, right panel). Global peak calling analysis showed comparable numbers of peaks in the CLIP and CLAP datasets, with the exception of PTBP1 replicate 2 (Fig. 3B). Notably, however, the authors’ replicate 2 lacked -tag controls (Fig. 3B). Thus, unlike for the PRC2 datasets, a determination of signal-to-noise levels for SAF-A and PTBP1 was not possible due to the absence of -tag controls. The missing data precludes a meaningful comparison of PRC2’s performance as an RBP relative to SAF-A and PTBP1.

**Figure 3.**
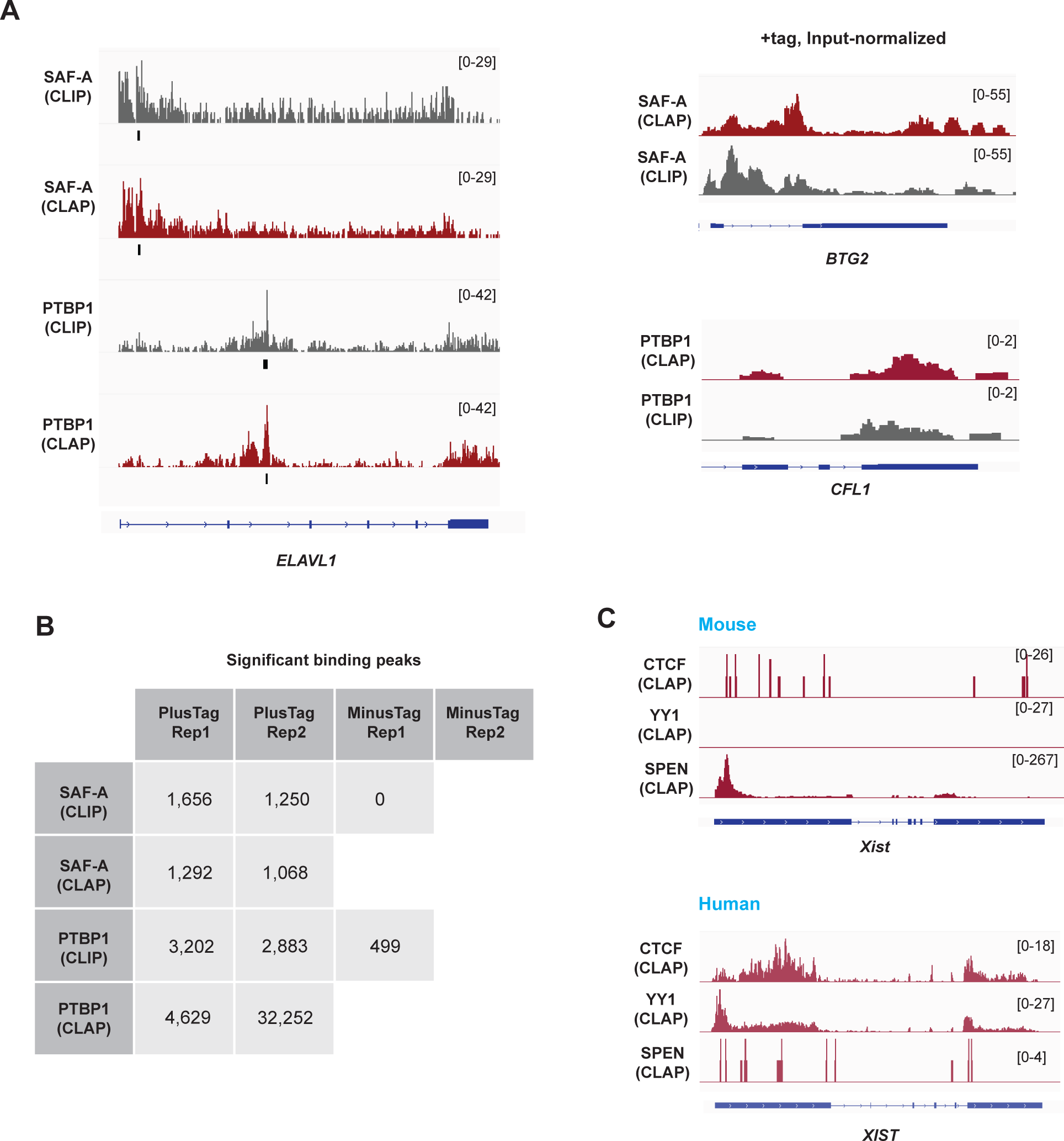
CLAP and CLIP both confirm that SAF-A and PTBP1 are RBP. (A) RNA-binding profiles of SAF-A and PTBP1 in +tag samples. Left: RPM-normalized RNA reads plotted across human ELAVL1 for CLIP and CLAP, with significant peaks marked by black bars. Right: Input-normalized enrichment profiles within BTG2 (top) and CFL1 (bottom) for CLIP and CLAP, showing no superiority of CLAP over CLIP. (B) Table summarizing significantly enriched over input CLIP and CLAP peak counts across datasets, with blanks indicating missing datasets in Guo et al.’s study. Note that absence of -tag controls for SAF-A CLAP and Replicate 2 (-tag) across all cases. (C) RNA-binding profiles of CTCF, YY1 and SPEN (RPM-normalized). CTCF and YY1 show strong signals across mouse Xist (top) and human XIST) (bottom) in +tag samples. In contrast, SPEN shows high signal at the 5’ end of mouse Xist, while signal across human XIST is minimal.

### Absence of input and -tag controls precludes analysis of SPEN, YY1, and CTCF

Similar criticisms can be applied to the analysis of SPEN as a positive control. There was only a single deposited mouse +tag SPEN CLAP sample. Because the study of Guo et al. was largely based on human +tag analysis, a +tag human sample should have been provided in order to compare RNA binding of SPEN to PRC2, PTBP1, and SAF-A. Moreover, a -tag sample should have been provided to assess the signal-to-noise difference. As is the case for PTBP1 and SAF-A, the missing data precludes a comparison of SPEN to PRC2 for their relative RNA binding potential. The authors therefore did not apply the same level of rigor to the analysis of SAF-A, PTBP1, and SPEN.

A similar critique applies to the authors’ analysis of the chromosome architecture protein CTCF and transcription factor YY1 ^25^. In these cases, the authors questioned the previously established RNA-binding status of CTCF and YY1 ^2,28,32–40^. Nevertheless, we were able to observe significant raw RNA signals from the human XIST and other transcriptomic regions in both CTCF and YY1 datasets (Fig. 3C). On the other hand, the lack of input control libraries for the CTCF and YY1 CLAP experiments prevented peak-calling and assessment of RNA binding significance. Additionally, the absence of -tag controls for both proteins precluded evaluation of the signal-to-noise ratio. Notably, the authors applied the -tag control to PRC2, PTBP1, and SAF-A to evaluate these proteins of interest as RBP. Consequently, no meaningful conclusions can be drawn from the CTCF and YY1 CLAP analyses conducted by Guo et al.

### Authors’ pipeline applied an arbitrary cutoff to PRC2 enrichment values

The substantial discrepancy between our findings and those of Guo et al. for PRC2 was surprising and could not be easily attributed to minor variations in bioinformatic methods. To ask if their near-complete lack of PRC2 RNA interactions could be attributed to a substantially different computational pipeline, we analyzed the same raw datasets using their reported pipeline (https://github.com/GuttmanLab/CLAPAnalysis). We plotted enrichment scores across the combined exon regions of XIST, comparing these values to those obtained using our own pipeline (Fig.1C). Interestingly, PTBP1 CLAP enrichment signals increased approximately 20-fold (from 3 to 60) over XIST, and SAF-A CLAP similarly showed a 20-fold increase (from 1 to 20)(Compare Fig. 4A, to Fig. 1C). The GFP CLAP negative control continued to show no enrichment signal over XIST, consistent with results from our independent pipeline (Fig. 1). This re-analysis demonstrates a close alignment of results with those presented by Guo et al. for PTBP1, SAF-A, and GFP (compare Fig. 4A to 1B, top panel).

**Figure 4.**
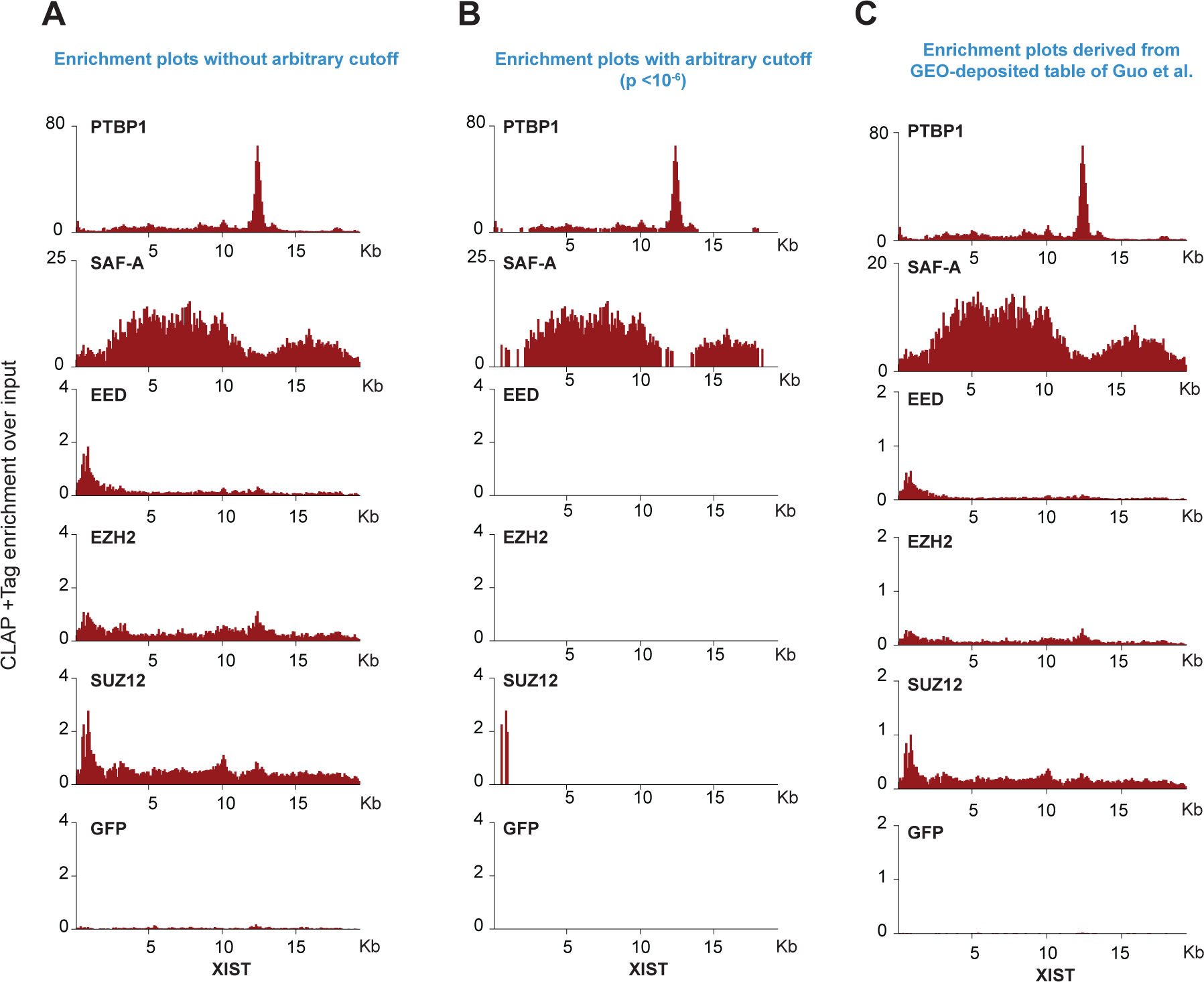
PRC2 qualifies as RBP when the pipeline of Guo et al. is applied to CLAP datasets. Input-normalized CLAP enrichment profiles for +tag human samples across XIST. All profiles were generated using Guo et al.’s computazonal pipeline (available on the authors’ GitHub) according to the authors’ guidelines. Combined XIST exons, excluding introns, are profiled. (A) Enrichment profiles were derived from raw sequencing files deposited by the authors on GEO, normalized by the the total library size (input) which includes both genome-mappable and unmappable reads. (B) Enrichment profiles using the same normalizazon method, but with exclusion of regions that did not meet an arbitrary p-value threshold of 10^−6^ (a “floor” for enrichment), as applied by Guo et al. (C) Enrichment profiles constructed from enrichment scores reported in the GEO-deposited table of Guo et al. without alterazon (GSE253466).

On the other hand, our re-analysis of PRC2 CLAP using the authors’ bioinformatic pipeline uncovered multiple unexpected findings. First, application of their method continued to show significant enrichment in XIST for all three PRC2 CLAP samples, well above the GFP background (Fig. 4A). This result stood in stark contrast to the complete absence of signal reported by Guo et al ^25^. (Fig 1B, upper panel). We therefore speculated that the authors might have applied an arbitrary cutoff. In their Methods section, Guo et al. stated a p-value cutoff of 10^−6^ for plotting and reporting purposes. We initially interpreted this as a cutoff applied exclusively to calling significant peaks. However, because of the large discrepancy between our enrichment plots and theirs remained, we surmised that this cutoff might have been applied additionally to the plots of enrichment values for XIST as well. Indeed, when we applied statistical significance threshold of p < 10^−6^ to enrichment scores, the XIST signals disappeared (Fig. 4B), and the enrichment plots now resembled their reported results ^25^ (Fig. 1B, upper panel). Thus, by applying an arbitrary “floor” to the enrichment plots, it appears that the authors bioinformatically discarded essentially all XIST signals in the PRC2 CLAP samples.

No explanation was provided for the use of the arbitrary cutoff. We questioned whether it was appropriate to apply it additionally when a significance threshold was already used in a prior step for calling CLAP peaks. To seek further clarification, we turned to the table of global CLAP enrichment scores deposited by the authors in GEO (GSE253466), reasoning that there should at least be a consistency between the GEO table and the plotted values (Fig. 1B, upper panel; Fig. 4C). Surprisingly, the authors’ own deposited table showed significantly enriched RNA signals for all three PRC2 subunits. Although the enrichment levels were lower than those of PTBP1 and SAF-A, they strongly contrasted with the absence of significant signals in the GFP negative control. This table-derived enrichment plot resembled the enrichment plots without a cutoff (compare Fig. 4A and 4C). We conclude that absent PRC2-XIST signals occurred only when the authors applied an arbitrary cutoff to enrichment scores, and the plotted “zero” values contradict the authors’ own reported data (GEO: GSE253466) showing positive enrichment scores in tabular format.

### Authors’ pipeline retained unmappable reads to obtain a normalization factor

By applying the authors’ bioinformatic pipeline, we uncovered additional unexpected findings. While PTBP1 and SAF-A signals *increased* approximately 20-fold over XIST relative to values reached by our independent pipeline, PRC2 CLAP signals *decreased* approximately 5-fold over XIST (Fig. 4A versus Fig. 1A). This was surprising, as we would normally expect enrichment factors to trend in the same direction if the experimental and bioinformatic pipelines were applied similarly across samples. This result led to us to consider differences in normalization factors.

Interestingly, Guo et al. attributed discrepancies in enrichment signals to our exclusion of stRNA reads (https://doi.org/10.1101/2024.11.02.621417) — i.e., the rRNA, 45S pre-rRNAs, tRNA, snoRNA, and snRNAs reads from non-unique RNAs that cannot be mapped to the genome. However, convention dictates that such repetitive RNAs be removed in preprocessing steps because of the repetitive nature and the inability to align the sequences to a unique genomic position. In addition, their repetitive nature prevents effective PCR de-duplication, making it impossible to distinguish unique sequencing reads from amplification artifacts. Consequently, including these RNAs in transcriptomic analysis can lead to inaccurate signal quantitation, significant skewing of results, and the creation of artifacts ^29,30,41^. Indeed, in the initial published account by Guo et al., the Methods section explicitly stated that stRNA reads were removed during preprocessing: “*PCR duplicates were removed using the Picard MarkDuplicates function*” and “*only reads that mapped uniquely in the genome and unambiguously to the human or mouse genomes were kept for further analysis”* ^25^. However, based on our re-analysis of their pipeline, this was apparently not what Guo et al. performed for CLAP analysis ^25^. Indeed, Guo et al. later revealed – contradicting their original publication – that these stRNA were retained (https://doi.org/10.1101/2024.11.02.621417). This practice of including unmappable RNAs not only contradicted convention, but also defied their earlier practice. For example, in the earlier SPEN/SHARP CLAP work by Jachowicz et al. ^42^, the Methods section stated that “*PCR duplicates were excluded from the analysis using the Picard MarkDuplicates function*”. The corresponding GEO record (GSE192574) also stated: “*Filtered paired-end reads were aligned using STAR aligner and uniquely mapped reads kept for analysis*”. As far as we could discern, retaining the unmappable reads in the work of Guo et al. ^25^ was neither reported in their Methods nor consistent with the laboratory’s standard practice.

Pertinent to the current investigation, inclusion of such unmappable reads could pose considerable normalization challenges, as their presence can create serious distortions and lead to inaccurate comparisons between samples. To determine the impact of including non-mappable reads on RNA enrichment calculations for the CLAP study, we calculated the ratio of unmappable reads to total reads in each +tag human CLAP sample. All three PRC2 subunits exhibited 10 to 20-fold higher levels of unmappable than mappable reads (Fig. 5A). The high number of unmappable reads was similar to that of the negative control, GFP. In contrast, unmappable reads accounted for only 30-40% of PTBP1 and SAF-A reads (Fig. 5A). These ratios might suggest that PRC2 is no more an RBP than GFP — as Guo et al. contended. However, when we examined fold-enrichment of mappable CLAP reads relative to mappable input, PRC2 showed greater enrichment than GFP (Fig. 5B, left panel). Thus, PRC2 CLAP samples showed enriched genome-mapped reads relative to the negative control.

**Figure 5.**
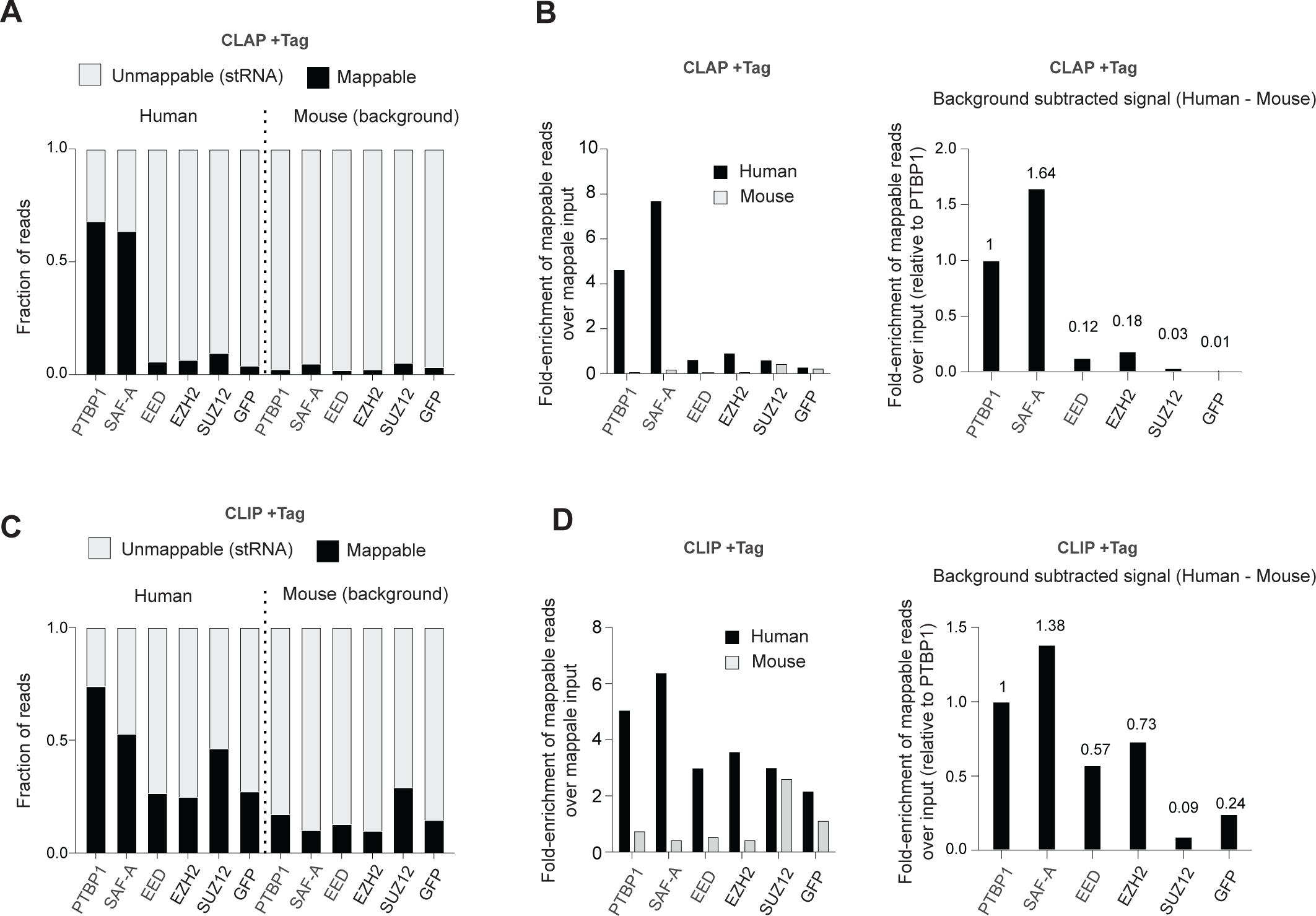
Levels of mappable and unmappable sequences in CLAP and CLIP. (A) Stacked bar plot showing the ratios of reads mapped to the reference genome. CLAP +Tag samples express Halo-tagged recombinant proteins in human cells mixed with wild-type mouse cells. Genome-mapped reads (black) and non-genome-mapped reads (e.g., rRNA, tRNA; gray) were calculated as a proportion of the total library reads. Mouse RNA serves as a background indicator. (B) Enrichment of genome-mapped reads in CLAP +tag samples. Fold enrichment in IP samples over input was calculated using genome-mappable ratios (left). Background-subtracted enrichment levels were determined by subtracting the mouse genome-mapped ratio, and normalizing to PTBP1 as a reference (right) (C) Ratios of genome-mapped reads in CLIP +Tag samples, represented as stacked bar plot similar to (A). (D) Enrichment of genome-mapped reads in CLIP +tag samples, represented as bar plots similar to (B).

### Inclusion of mouse reads inflated human PRC2 CLAP background

Still, the degree of PRC2 enrichment over input was only a fraction of that observed for PTBP1 and SAF-A (Fig. 5B, left panel). To investigate why the authors’ PRC2 CLAP suffered from such high background, we delved further into the nature of the unmappable reads. Per the authors’ experimental design, *human* cells expressing Halo-tag proteins were mixed with wild-type *mouse* cells to enable examination of reassociation artifacts. Scrutiny of their pipeline revealed that Guo et al. included the <u>mouse</u> reads in the analyses of <u>human</u> CLAP and CLIP ^25^. As discussed above, inclusion of unmappable reads, especially from a foreign species, may cause unintended distortions.

We re-analyzed their data by subtracting the expected background level due to mouse reads in +tag experiments. Intriguingly, the +tag human CLAP for EED, EZH2, and SUZ12 subunits exhibited higher signals (0.12, 0.18, and 0.03, respectively) than fully depleted GFP signals (0.01) (Fig. 5B, right panel). Among PRC2 subunits, EZH2 demonstrated the strongest enrichment, with an 18-fold enrichment over GFP, whereas SUZ12 showed a 3-fold enrichment (Fig. 5B, right panel). These results align with UV RNA-protein crosslinking studies showing higher crosslinking frequencies in EED and EZH2 surface residues compared to SUZ12 ^14^. Our re-analysis using the authors’ pipeline therefore agrees with the analysis using our independent method (Fig. 1), with both validating PRC2 as an RBP.

However, even after subtracting the background level when using the pipeline of Guo et al., PTBP1 and SAF-A CLAP signals were 5 to 15 times more enriched than those of PRC2 subunits (Fig. 5B, right panel). This result did not agree with findings from our independent method (Figure 1). To investigate further, we turned to the authors’ conventional <u>CLIP</u> datasets. Without subtracting background, human PRC2 CLIP datasets also showed more unmappable reads than SAF-A and PTBP1 (Fig. 5C). However, the mappable signals for EED and EZH2 CLIP remained substantially greater (Fig. 5D, left panel), as was the case for CLAP (Fig. 5B, left panel). Notably, the background mouse reads in the human SUZ12 and GFP CLIP samples were 7- and 3-times higher, respectively (Fig. 5D, left panel). When the mouse background was appropriately subtracted, EED and EZH2 signals became again greater than SUZ12 and GFP (Fig. 5D, right panel). The CLIP results therefore also validate PRC2 as an RBP, consistent with the analysis provided by our independent method (Figure 1).

### Selective retention of read duplicates further inflated the background

Given the high stringency of denaturing washes in CLAP, the large unmappable (stRNA) fractions for PRC2 were surprising to us. Moreover, for PRC2 and GFP CLAP, enrichment of mappable reads in the IP was lower than that for input (Fig. 5B, left panel, fold-enrichment <1). This was surprising, because if PRC2 were not an RBP and its signals derived strictly from the background, PRC2 signal enrichment should be similar to the input “background” (Fig. 6A, bottom panel “Non-RBP”; 6B), whereas a true RBP would yield signals in the IP-captured fraction in significant excess over the input background (Fig. 6A, top panel “True RBP”; 6B). For a deeper analysis, we compared levels of unmappable stRNA for CLAP versus CLIP and for human versus mouse (Fig. 6C). When comparing percentages of unmappable stRNA in various <u>inputs</u>, we found similar levels of stRNA in all samples (PTBP1 and SAF-A versus PRC2 components). However, in CLAP and CLIP samples, stRNA levels were 2- to 3-fold higher for PRC2 than for PTBP1 and SAF-A. Curiously, this discrepancy was observed only in the proportions of human reads of IP samples, not in the mouse reads, which serve as the background from these samples (Fig. 6C). This great over-representation of stRNA in the human reads of PRC2 CLAP would have further diluted the specific signals and would have therefore been another major factor contributing to extremely low PRC2-RNA signals in the +Tag (human) CLAP, aiding the authors’ argument that PRC2 is not an RBP.

**Figure 6.**
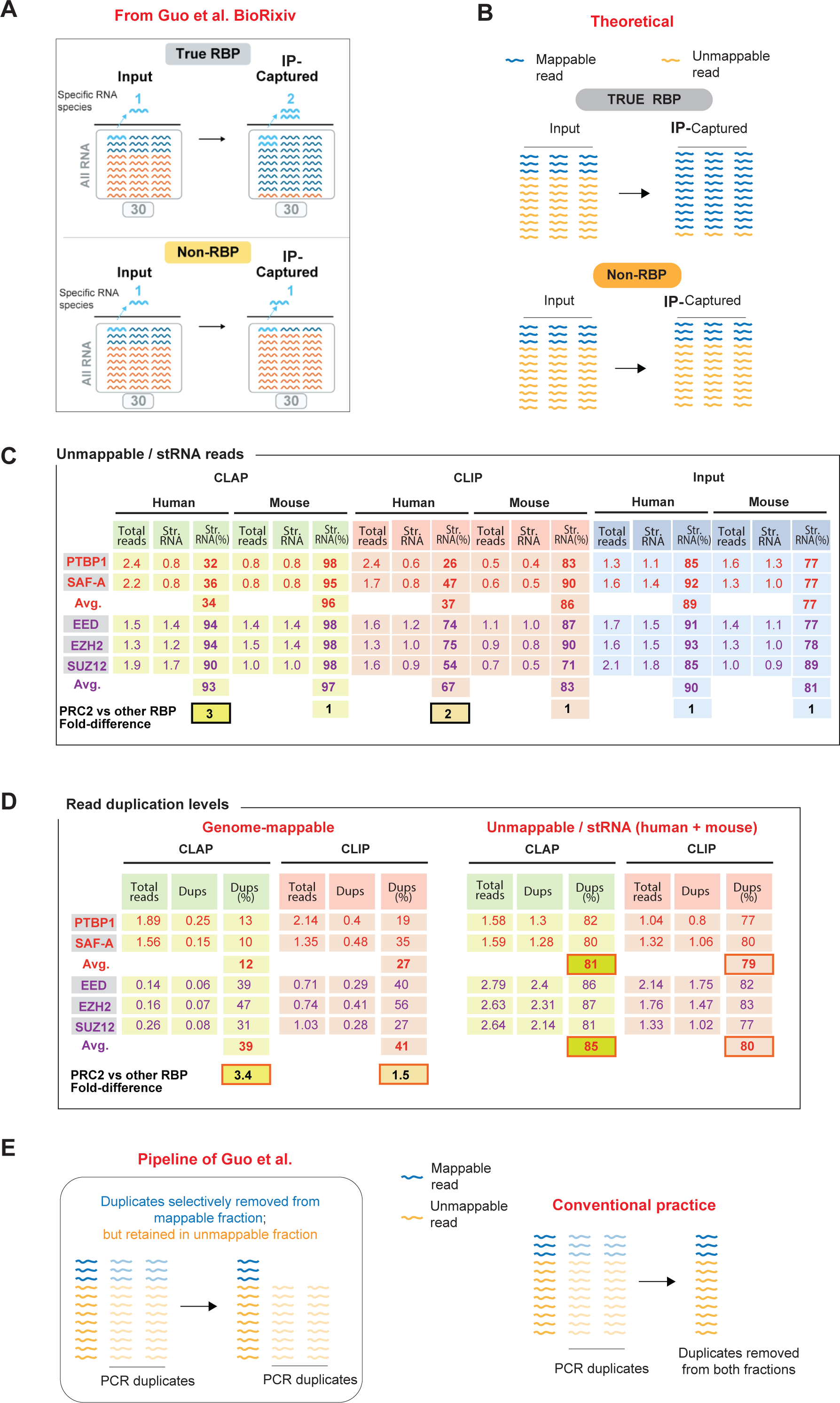
Retention of unmappable reads and selective removal of read duplications from the mappable fraction resulted in severe deflation of PRC2 enrichment. (A) Schematic representation of the expected genome-mapped reads enrichment in RBPs and non-RBPs. Left panel is reproduced without alteration from Guo et al. (B) Right panel illustrates the anticipated enrichment pattern in non-RBPs, where the background matches the enrichment level observed in the input. (C) Summary of unmappable reads (stRNA) in CLAP and CLIP experiments. Total read counts per transcriptome and read counts for stRNA reads are shown in millions of reads for each column, with %stRNA shown in parallel. Mouse and human reads shown separately. CLAP and CLIP experiments for each species are shown side by side with Input for comparison. Average (Avg) stRNA % shown for PTBP1/SAF-A together (red font), or PRC2 subunits together (purple font). Fold-difference between these averages is shown at the bottom in black boxes. (D) Summary of mappable (unique sequences) versus unmappable (stRNA) read duplications in CLAP and CLIP experiments. Per pipeline of Guo et al., the unmappable fraction is a composite of mouse + human reads. Total reads and the number of read duplications (Dups) per transcriptome are shown in millions of reads, with % duplication shown in parallel. CLAP and CLIP experiments are shown side by side for mappable versus unmappable fractions. Average (Avg) % shown for PTBP1/SAF-A together (red font), or PRC2 subunits together (purple font). Fold-difference between these averages is shown at the bottom in black boxes. (E) Selective removal of duplicates from the mappable fraction. Duplicates are retained in the stRNA fraction. Left panel highlights the issue of exclusively removing duplicate reads from genome-mapped reads, stRNA reads remain unadjusted. Right panel shows the conventional practice of applying duplicate removal to both genome-mapped and unmappable reads.

Read duplicates can arise either from cloning of two distinct RNA molecules or from PCR over-amplification of a target. Conventional practice generally recommends removal of read duplicates in preprocessing steps prior to analysis ^29,30,41^, as their inclusion of duplicates can artificially inflate transcript counts and distortion of RNA enrichment calculations, especially when PCR amplification cycles are high. Guo et al. acknowledged this practice, as their Methods section stated: “*PCR duplicates were removed using the Picard MarkDuplicates function*” ^25^. However, based on both their bioRxiv communication (https://doi.org/10.1101/2024.11.02.621417) and computational pipeline (https://github.com/GuttmanLab/CLAPAnalysis), they did not seem to have implemented this practice and instead included stRNA reads and their PCR duplicates in the analysis. Notably, due to their repetitive nature, PCR duplicates generated by stRNAs cannot be reliably removed even with advanced tools such as *Picard* or *Sambamba*. If the authors wished to include duplicates, an effective way to account for PCR duplicates and obtain accurate read counts would be to incorporate Unique Molecular Identifiers (UMIs) in the experiment. UMIs help mitigate this issue by distinguishing original RNA molecules from PCR duplicates, enabling precise transcript quantitation and reducing amplification bias. To our knowledge, Guo et al. did not incorporate UMIs in their study and instead based their enrichment calculations on a practice that has long been recognized as inadequate.

As noted above, PRC2 CLAP samples contained three times more stRNA reads than the PTBP1 and SAF-A samples (Fig. 6C). Given that Guo et al.’s pipeline did not remove PCR duplicates from stRNAs, we questioned whether the increased stRNA read proportions could have resulted from accumulation of PCR duplicates during the generation of PRC2 CLAP and CLIP samples. Since both the mappable (e.g. mRNA) and unmappable (stRNA) fractions were derived from the same transcriptome and were amplified in the same PCR reaction, the degree of read duplication must be similar across both fractions for each CLAP and CLIP samples. To calculate the percentage of PCR duplicates, we used the *Picard* tool. However, as expected, due to the repetitive nature of stRNA, removal of duplicates proved challenging as the duplicates could result from limited sequence complexity of repetitive elements. This resulted in similar proportions of PCR duplicates between PRC2 samples and PTBP1 and SAF-A (Fig. 6D, right panel, human + mouse composite shown). We then examined duplication levels in the mappable fraction (Fig. 6D, left panel). Strikingly, read duplicate levels were 3.4 times higher for PRC2 than for SAF-A and PTBP1. The discrepancy for PRC2 was greater for CLAP than for CLIP, but both sets of experiments showed a strong bias towards PRC2 samples — in contrast to those for PTBP1 and SAF-A.

A close examination of the computational script of Guo et al. revealed that it selectively removed duplicates only from the mappable fraction, while retaining them in the stRNA fraction (Fig.6E). In order to achieve this, Guo et al. initially separated the unmappable from mappable reads, performed de-duplication selectively on the mappable reads, and then merged the separately treated mappable and unmappable reads (Fig. 7A). No explanation was provided for these unusual steps, nor was the selective removal documented in the Methods (that we could find). In fact, these steps directly contradicted their explicit statement in the Methods that only uniquely aligned and de-duplicated reads were used in their analysis ^25^. The fact that PRC2 CLAP and CLIP samples contained a substantial fraction of PCR duplicates far exceeding that of the PTBP1 and SAF-A samples (Fig. 6C, 6D) may result from several possibilities, including stochastic variation or introduction of additional PCR cycles for PRC2 during library generation, for example. Regardless of the cause, it would have been appropriate to apply a computational pipeline that mitigates technical constraints to avoid enrichment distortions that would disproportionately affect one sample over another due to a high stRNA content.

**Figure 7.**
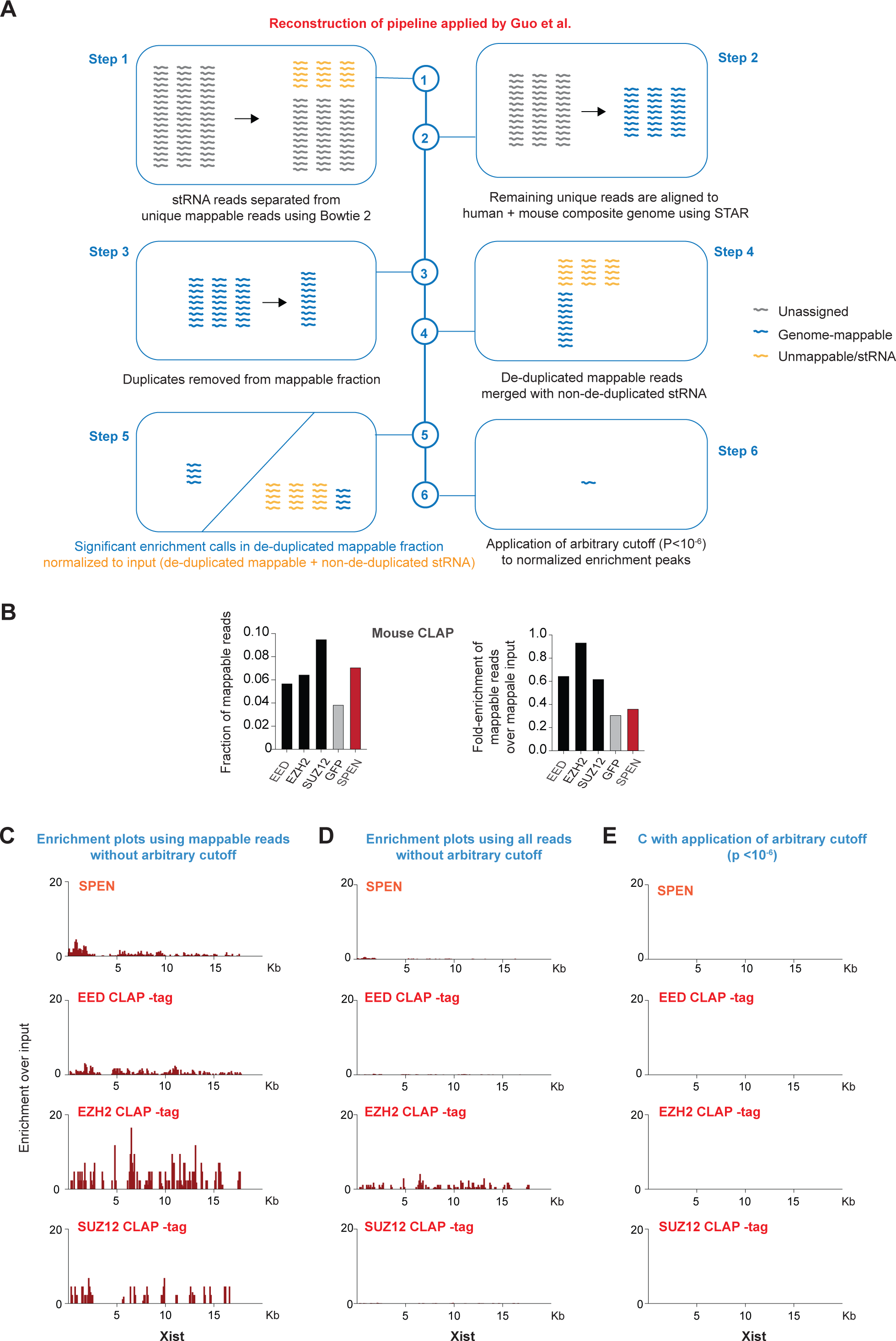
Summary of authors’ pipeline and application to SPEN. (A) Reconstruction and summary of the pipeline utilized by Guo et al. In Step 1, CLIP and CLAP reads are aligned to the structural RNA genome using Bowtie2 and stRNA reads are filtered away for Steps 2-3. In Step 2, the remaining “unique” reads are mapped to the mouse (mm10) and human (hg38) as a composite genome using STAR aligner. In Step 3, duplicate reads are removed from the genome-mapped reads only. In Step 4, the de-duplicated mappable fraction is re-united with non-deduplicated stRNA reads. In Step 5, global enrichment calculations are performed on the merged dataset. In Step 6, an additional and arbitrary cutoff is applied to the enrichment set. (B) Left panel: Ratio of genome-mapped reads in for mouse PRC2 subunits, GFP, and mouse SPEN/SHARP (GSE192574) CLAP samples., Right panel: Fold-enrichment of mappable reads relative to mappable input for PRC2 subunits, GFP, and SPEN. **(C-E)** Mouse SPEN/SHARP (GSE192574) and PRC2-tag CLAP (mouse) results are shown using different criteria for Xist RNA binding. (C) Enrichment plots using mappable reads without a cutoff. (D) Enrichment plots shown using all reads (mappable + unmappable) without an arbitrary cutoff. (E) Enrichment scores calculated using all reads (mappable + unmappable) with an arbitrary cutoff of P<10^−6^.

### CLAP reaffirms PRC2 and SPEN as robust RBPs

As summarized in Figure 7A, Guo et al. applied a highly unconventional normalization pipeline in that they (i) initially filtered away the stRNA reads (Step1) and mapped the remaining unique reads to the genome (Step 2), (ii) removed read duplicates selectively from the mappable reads fraction (Step 3), (iii) then re-introduced the stRNA fraction without de-duplicating the stRNA (Step 4, merge), (iv) retained foreign species (mouse) RNA reads (Step 3), thereby adding to the background, (iv) performed enrichment analysis for significant peaks (Step 5), and (v) applied a 2^nd^ enrichment cutoff to define an arbitrary “floor” (Step 6; Fig. 4B). This unusual pipeline fully accounts for the discrepancy between our observations and those of Guo. Notably, even with the excess stRNA reads in PRC2 CLAP and selective duplicate removal in the mappable fraction, PRC2 still demonstrated significant RNA enrichment (Fig. 4A,C). <u>Only by applying</u> <u>the unjustified enrichment cutoff on top of the selective duplicate removal were the authors able to</u> <u>eliminate PRC2 as an RBP</u> (Fig. 4B). The authors’ unconventional practices, in this case, had the specific consequence of artificially deflating PRC2 CLAP signals and inflating PRC2 CLAP background — thereby leading to the impression that PRC2 is not a robust RBP.

Finally, we asked how SPEN (a.k.a. SHARP) would fare if the pipeline were applied to the authors’ previously published mouse CLAP study ^42^. SPEN is a known RBP with affinity for mouse Xist’s Repeat A region ^42–45^. Here we re-analyzed the Guttman Lab mouse SPEN dataset ^42^ using the current pipeline of Guo et al. ^25^ and compared the output to those of the “CLAP minus tag” datasets in which human cells are untagged and the admixed mouse cells are PRC2-halo-tagged (thereby enabling mouse CLAP analysis) ^25^. The number of total mappable reads was similar between SPEN and PRC2 (Fig. 7B, left panel). However, overall enrichment was somewhat greater for PRC2 than for SPEN; in fact, the SPEN enrichment was similar to that of GFP’s (Fig. 7B, right panel). When enrichment values were plotted across Xist RNA using only mappable reads (per convention and reported Method of Jachowicz et al. ^42^), SPEN enrichment was similar to that of PRC2’s EED subunit (Fig. 7C). However, when plotted using all reads (mappable + stRNA), SPEN performed similarly relative to PRC2 (Fig. 7D). Lastly, when the same arbitrary enrichment cutoff (*P* < 10^−6^) was applied to all samples, all signals disappeared (Fig. 7E). <u>Thus, PRC2 and SPEN were treated</u> <u>differently. If similar criteria were used for PRC2 and SPEN, SPEN would also be excluded as an Xist binder</u> <u>and would not meet their definition of an RBP.</u> In the published work of Jachowicz et al., the Guttman group followed convention and removed stRNA reads and duplicates. It is unclear why Guo et al. disregarded this standard practice and retained stRNA and duplicates in the PRC2 CLAP study.

In summary, using two independent methods (including the authors’ own pipeline), our re-analysis of the CLIP and CLAP datasets deposited in GEO failed to reproduce the presentation and the conclusions of Guo et al. ^25^ Specifically, our examination revealed significant PRC2 enrichment across the transcriptome, including XIST RNA. Discrepancies can be attributed to reporting deficiencies, selective filtering, and differential treatment of RBPs. The net effect is that the pipeline inflated PRC2’s background reads and suppressed the mappable IP fraction, creating the misleading impression that PRC2 is not a robust RBP. If the pipeline were applied to the authors’ own SPEN/SHARP CLAP data, this well-established RNA-binding protein would also fail the definition of an RBP. A large body of work currently supports functional PRC2-RNA interactions – such as in the fine-tuning of gene expression via transcriptional pausing ^7,22^, targeting to specific chromatin sites ^1,5,26,46^, control of a protein’s catalytic activity ^14,16,19^, regulation of PRC2 eviction from RNA versus DNA ^1,4,17,19,47^, and control of SINE B2/ALU ribozyme activity during the stress response ^23^ ^24^. In addition, a recent pre-print has argued that amino acid residues involved in PRC2-RNA interactions are inefficiently UV-cross-linked to RNA, a fact that may account for reduced signals for PRC2 relative to other RBPs^48^. We conclude that the CLAP analysis provided by Guo et al. does not invalidate established biochemical, epigenomic, and functional studies for PRC2-RNA interactions. On the contrary, our independent analysis of their dataset reaffirms PRC2 as an RBP.

## METHODS

### Datasets

All CLIP and CLAP datasets were retrieved from GEO: GSE253477 and GSE192574 ^25,42^.

### Read sequence processing and alignment to the reference genome

FastQ files were preprocessed by performing adapter trimming using Trim Galore. To exclude reads mapping to non-target regions, the resulting trimmed reads were initially aligned using STAR aligner ^49^ to a mouse (mm10) reference composite comprising genomic sequences from rRNAs, small nuclear RNAs (snRNAs), small nucleolar RNAs (snoRNAs), 45S pre-rRNA, and tRNA. Unaligned reads were further filtered by aligning them to a human (hg38) reference composite containing the corresponding human-specific non-target genomic elements. Subsequently, the remaining reads were aligned to a composite reference including both mouse (mm10) and human (hg38) genomes. Duplicate reads were removed using sambamba_v0.6.6 ^50^. BAM files were sorted, indexed, and split into forward and reverse strands. Strand-specific coverage tracks were generated by processing BAM files with the bamCoverage tool from deepTools ^51^, using a bin size of 10 and normalizing with RPKM (Reads Per Kilobase Million) to account for differences in library size and gene length.

### Normalization by fold-enrichment relative to input

FastQ files of IP libraries (CLIP or CLAP) and their corresponding Input samples were preprocessed and aligned as described earlier. When two replicates of the same type were available, they were merged, preprocessed, and aligned. Reads were then separated into Forward and Reverse strands. To ensure comparability between the IP and its corresponding Input, the larger library (either IP or Input) was randomly downsampled to match the size of the smaller library, with this process being performed separately for Forward and Reverse strands. The downsampled library pairs (IP and Input) were then used as input for MACS2 analysis ^52,53^ with the parameter ‘--format BAM –nomodel –bdg – SPMR’ to generate signal tracks (.bdg files) for both IP and Input libraries. MACS2’s ‘bdgcmp’ utility was employed with the Fold Enrichment (FE) model to compare the IP track to its corresponding Input track. This comparison provided the fold enrichment signal of the IP over the Input, effectively normalizing the IP signal to the input control.

### Quantification for CLIP and CLAP signal enrichment over genes

FDR-corrected q-values and enrichment over input metrics for each gene were calculated using Cufflinks v2.21, with the strand-specific library flag (-library-type fr-firststrand) applied. Sorted BAM files, prior to strand splitting, were provided to the cuffdiff module of the Cufflinks program. Enrichment and statistical analyses were conducted on the annotated genes from the GRCh38 (human) or GENCODE version M25 (mouse) reference gene assemblies. Scatter plots were generated using GraphPad Prism. The y-axis represents CLIP and CLAP signals on the genes normalized by the input signal, and then the dots were rearranged with the input abundance (log2 FPKM) on the x-axis for visualization.

### Peak-Calling

Strand-separated BAM files were used for peak calling, and significantly enriched peaks over input reads were indicated by MACS2 ^52,53^ program with the no-shifting model parameters (-p 0.01 -f BAM -- nomodel -g hs).

### Motif analysis

HOMER ^54^ module (findMotifs.pl) analyzed enriched RNA motifs from the sequences of significant peak regions. Significant peak regions commonly observed in biological replicates that do not overlap with negative control (minus tag) peaks are used for the analysis.

### CLAP and CLIP Analysis following Guo et al’s computational pipeline

All the CLAP and CLIP IP libraries were processed following the instructions from Guo et al. GitHub repository (2024) (https://github.com/GuttmanLab/CLAPAnalysis). Paired-end FASTQ reads were adapter-trimmed using Trim Galore!, then aligned to a species specific genome reference containing repetitive and structural RNAs (e.g., ribosomal RNAs, snRNAs, snoRNAs, 45S pre-rRNAs, tRNAs) using Bowtie2. Mapped reads were retained for further analysis, while unmapped reads were converted to FASTQ files and aligning to the combined genome reference containing the mouse (mm10) and human (hg38) genomes using STAR aligner. PCR duplicates from STAR-aligned reads were removed using Picard’s MarkDuplicates utility. Reads were further processed to exclude blacklisted repeat regions, using service blacklist files from the GitHub repository. Processed reads were then merged with the Bowtie2-mapped repetitive and structural RNA reads, creating the final alignment file for downstream analysis. Input library FASTQ files were processed following the same abovementioned steps, with the exception that they did not undergo PCR duplicate removal at any step. The final alignment files for IP and corresponding Input libraries were used to calculated genome-wide enrichment using the ‘Enrichment.jar’ script, along with the gene annotation file provided in the GitHub repository. Default parameters, including 1000 bp window size, were applied for this analysis. Along with the default parameters, the enrichment calculation step also takes the number of reads for IP and Input libraries for scaling. To get the number of mapped reads from the final merged alignment files, we used samtools view utility (with parameter ‘-c’).For the analysis results involving only genome mapped reads, samtools view utility was used with reference genome chromosome names (e.g, hg38: chr1-22, X, Y and M; mm10: chr1-19, X, Y and M) to extract reads assigned specifically to those chromosomes. For species-specific experiments, we used hg38 genome for human samples and the mm10 genome for mouse samples. Mixed-species experiments utilized a combined hg38/mm10 genome to ensure comprehensive alignment and enrichment calculation.

## ACKNOWLEDGMENTS

We thank Richard Jenner, Roberto Bonasio, Rodrigo Aguilar, and the Lee Lab for discussion and feedback. This work was supported by an NIH grant (R01-HD097665) to JTL.

